# Targeted genomic screen reveals focal long non-coding RNA copy number alterations in cancer

**DOI:** 10.1101/113316

**Authors:** Pieter-Jan Volders, Jo Vandesompele, Steve Lefever, Shalina Baute, Justine Nuytens, Katrien Vanderheyden, Björn Menten, Pieter Mestdagh

## Abstract

The landscape of somatic copy-number alterations (SCNAs) affecting long non-coding RNAs (lncRNAs) in human cancer remains largely unexplored. While the majority of lncRNAs remains to be functionally characterized, several have been implicated in cancer development and metastasis. Considering the plethora of lncRNAs genes that is currently reported, it is conceivable that several lncRNAs might function as oncogenes or tumor suppressor genes.

We devised a strategy to detect focal lncRNA SCNAs using a custom DNA microarray platform probing 20 418 lncRNA genes. By screening a panel of 80 cancer cell lines, we detected numerous focal aberrations targeting one or multiple lncRNAs without affecting neighboring protein-coding genes. These focal aberrations are highly suggestive for a tumor suppressive or oncogenic role of the targeted lncRNA gene. Although functional validation remains an essential step in the further characterization of the involved candidate cancer lncRNAs, our results provide a direct way of prioritizing candidate lncRNAs involved in cancer pathogenesis.

## Introduction

The cancer genome is marked by large numbers of genetic and non-genetic alterations. The greater majority of those are somatic. Only a small fraction of the somatic mutations, the so-called driver mutations, contribute to cancer development by activating or inactivating specific cancer genes. The remainder are passenger mutations that do not confer growth advantage but were acquired at some point during cancer cell proliferation(1). Differentiating between driver and passenger mutations is one of the biggest challenges in the quest for new cancer genes and putative therapeutic targets. While somatic alterations can be as small as a single nucleotide substitution, insertion or deletion, somatic copy-number alterations (SCNA) affect the largest fraction of the genome(2). In some cases, SCNA affect entire or partial chromosome arms. The ability to detect these genetic/genomic alterations using (molecular) cytogenetic methods has made large SCNA historically the best studied cancer associated genetic alterations. Many well-known oncogenes and tumor suppressor genes have been initially identified as targets of recurrent genomic amplifications or deletions, respectively. Notable examples are tumor suppressor genes PTEN(3) and RB1(4) and oncogenes HER2 (ERBB2)(5) and the MYC-family of transcription factors(6,7). The resulting diagnostic and therapeutic successes have made cancer SCNA subject of many studies. Additionally, the advent of array comparative genome hybridization (array CGH) platforms that enable robust identification of small SCNAs greatly improved our knowledge of the cancer genome(8–10).

As cancer genetics until now mainly focused on protein-coding genes, not much is known on SCNAs affecting non-coding RNA genes in cancer. In recent years, our knowledge on the non-coding genome has expanded enormously. This is especially the case for the class of long non-coding RNAs (lncRNAs), consisting of genes with transcripts larger than 200 nucleotides that do not encode proteins. In the past 5 years, ten thousands of human lncRNAs have been reported and catalogued, making this the largest genetic class in the human genome(11). While the bulk of lncRNAs remains to be functionally annotated, they have been implicated in many important normal cellular processes such as dosage compensation(12), chromatin remodeling(13), and cell differentiation(14); when deregulated, they play a role in disease as well, including cancer(15).

The discovery of cancer associated lncRNAs such as HOTAIR(16), MALAT1(17) and PVT1(18) uncovered an important role for lncRNAs in oncogenesis. The reason for the current hiatus in our knowledge on lncRNA SCNAs is the fact that the majority of lncRNA annotations are very recent. Most commercially available platforms are based on older genomic annotations (with no probes for lncRNAs, or probes for as yet unannotated lncRNAs) or lncRNAs are simply overlooked in the data analysis. Indeed, recurrent SCNAs outside of protein coding regions have been reported(2,19). To overcome this problem, existing DNA microarray platforms have been repurposed and probe content was reannotated with current lncRNA annotation(20,21). One such effort resulted in the discovery of the oncogenic FAL1 (focally amplified lncRNA on chromosome 1) lncRNA in ovarian cancer(21). While the potential of this approach lies in its ability to make use of the large amount of publically available DNA microarray data, the used platforms have several disadvantages for the discovery of putative cancer associated lncRNAs. Whole cancer genome sequencing has the potential in principle to circumvent these limitations, but the method is still relatively expensive, and challenging in terms of data-analysis. Consequently, public databases (e.g. TCGA) are mainly populated with targeted exome sequencing datasets, again focusing on protein coding genes.

Here we present a targeted and cost-effective approach to identify focal lncRNA SCNA based on a custom DNA microarray covering 20 418 lncRNA transcripts and their flanking protein coding genes. We show the ability of this platform to detect focal aberrations that only affect lncRNA exons and not encompass their flanking protein coding genes. By analyzing the DNA of 80 cancer cell lines covering 11 cancer subtypes we reveal that lncRNAs are frequently targeted by focal aberrations in human cancer. In addition, we have generated a dataset with putative oncogenic and tumor suppressor lncRNAs for future functional studies.

## Results

### A targeted platform to detect focal copy number changes in lncRNA genes

LncRNAs are underrepresented on commercial array CGH platforms and the mean chromosomal distance between the probes on these arrays makes them unsuitable to detect small aberrations that only involve (part of) a single lncRNA gene (Figure S1, Supplementary material).

In order to detect small and focal SCNAs that only affect lncRNA exons, we designed a custom 180k CGH array covering intergenic lncRNA exons and the nearest exons of their flanking protein coding genes. To this purpose, we constructed a database with 52 324 non-redundant exons derived from all transcripts listed in LNCipedia (Figure 1, Figure S2 and Figure S3). The database was subsequently extended with protein coding gene annotation from Ensembl. Next, we designed probes using the genomic sequence of the lncRNA exons and the two nearest exons of the flanking protein coding genes. By removing duplicate probes in overlapping exons and selecting additional probes for transcripts with fewer exons, we were able to cover the majority (94%) of the transcripts with at least 10 probes (Figure S4). Only 1.2% of lncRNAs could not be covered by any probe. For 95% of the lncRNA transcripts we succeeded in designing 2 probes for each flanking protein coding exon.

**Figure 1.**
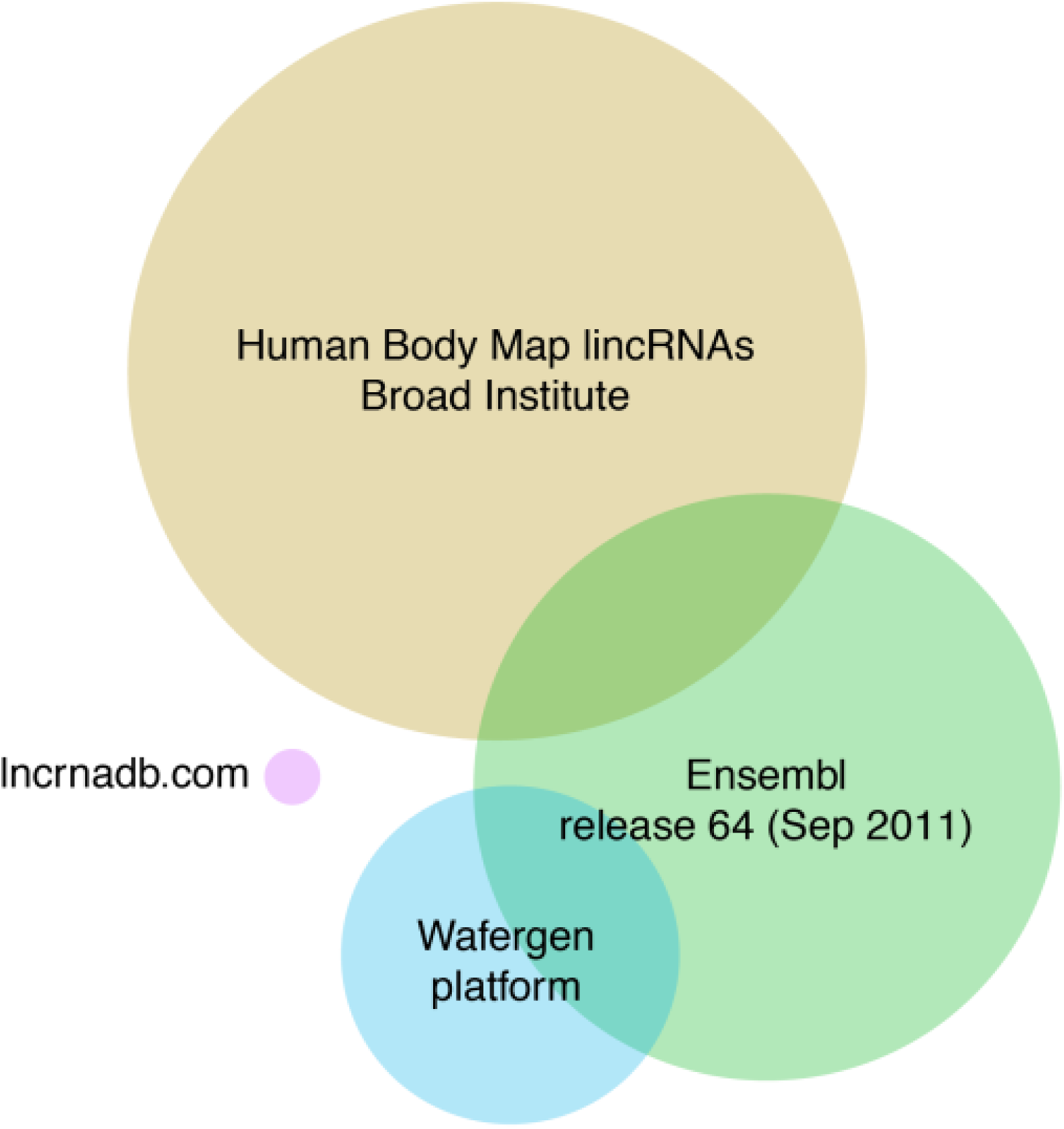
Euler diagram of the different IncRNA sources. The circle diameter and overlap correspond to the number of lncRNAs. The sources include several lncRNA databases and the lncRNAs on the Wafergen SmartChip Human LncRNA1 Panel (http://www.wafergen.com/products/smartchip-panels/smartchip-human-lncrna1-panel/)

To assess the quality of our custom array CGH platform, we compared the profiles for 60 cancer cell lines (NCI-60 subset) to publically available profiles of two different array CGH platforms. The average log ratio in 1 Mb bins was calculated and correlated between the different platforms. These correlations were compared with correlations among unrelated cell lines (Figure S5 and Figure S6). Correlation between the same cell lines across different platforms was high (median Pearson’s correlation = 0.70), validating the quality of our profiles. As expected, cell lines derived from the same individual (such as NCI/ADR-RES and OVCAR-8) are also highly correlated (Pearson’s correlation = 0.74). In addition, this analysis revealed problems with 2 DNA samples (HCT-15 and CAKI-1) as the obtained profiles showed poor correlation with publically available profiles. This poor correlation remained unresolved by repeating the hybridization. As such, results from these two cell lines should be interpreted with care.

### Frequent focal lncRNA copy number alterations in cancer cell lines

To explore focal lncRNA SCNAs in cancer, we analyzed DNA from 80 cancer cell lines covering 11 cancer subtypes with our custom DNA microarray (Table 1). An extensive filtering was performed on the resulting segments to shortlist focal lncRNA SCNA alterations. To be considered a lncRNA SCNA, a segment should (1) overlap with exonic lncRNA sequence, (2) not be contained within known segmental duplications, (3) overlap with at most 3 known variants and (4) have an absolute average log-ratio that is larger than 1.5 (reflecting homozygous deletions and gene amplifications). In the case of an amplification, an additional requirement was that the segment includes the entire transcript. Finally, to withhold a focal SCNA (5), the segment cannot overlap any of the flanking protein coding gene exons. Using these settings, 173 focal SCNAs affecting 136 lncRNAs in at least one cell line were identified (Figure 2, Supplementary Table 1). The majority of these lncRNAs (111) is affected in a single cell line, 16 are affected in 2 cell lines, 7 in 3 cell lines, 1 in 4 cell lines and 1 in 5 cell lines. By confining the relative difference in log-ratio between the segment covering the lncRNA and the segment covering the flanking protein coding genes, it is possible to retain superimposed SCNAs (for instance a large hemizygous deletion that contains a smaller homozygous deletion). A more stringent subset of 76 lncRNA SCNA is obtained if we require that the flanking protein coding gene does not show any copy number change (Figure S7, Supplementary Table 2).

**Table 1.**
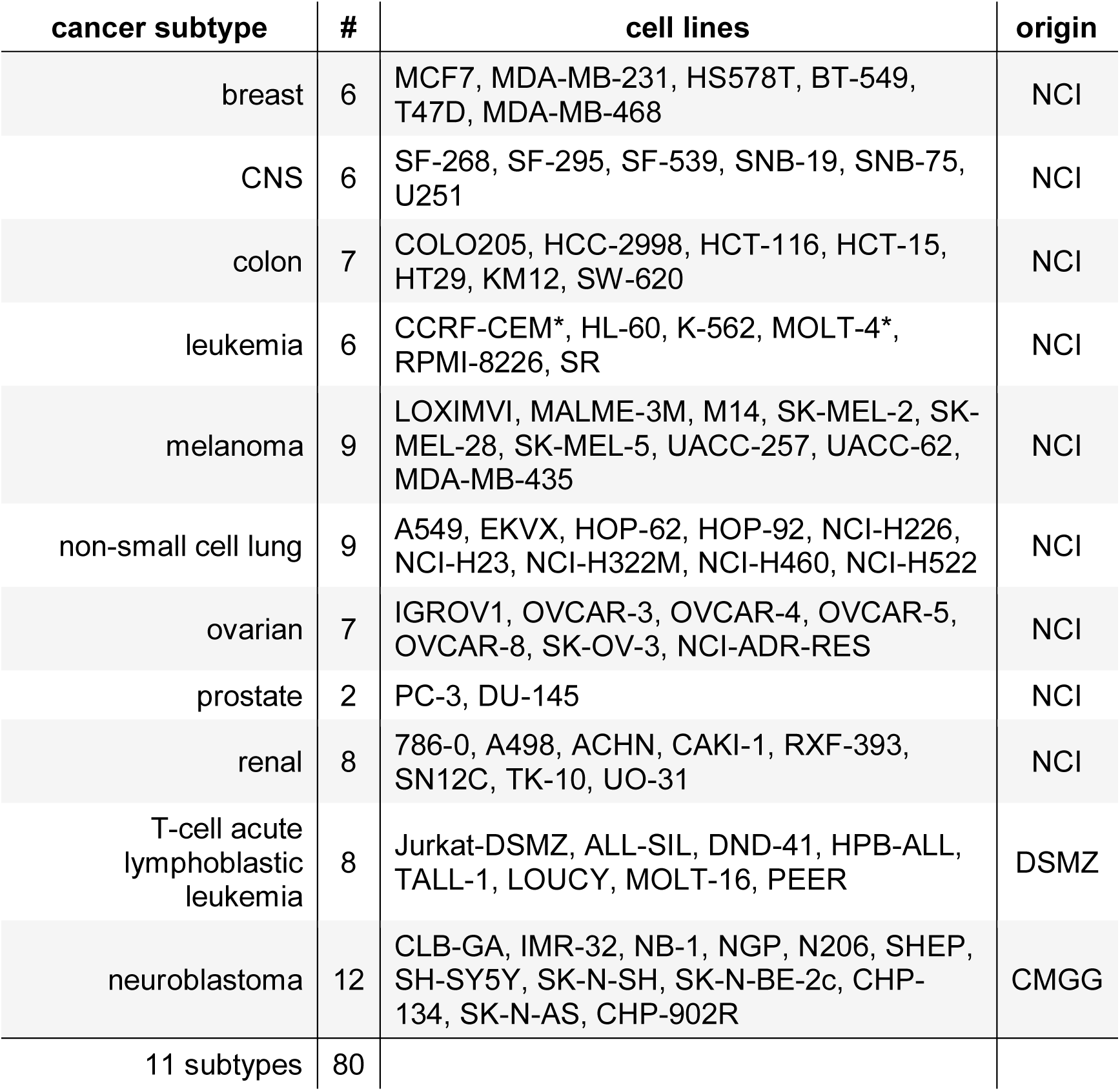
Overview of cell line panel and cell line origin

**Figure 2.**
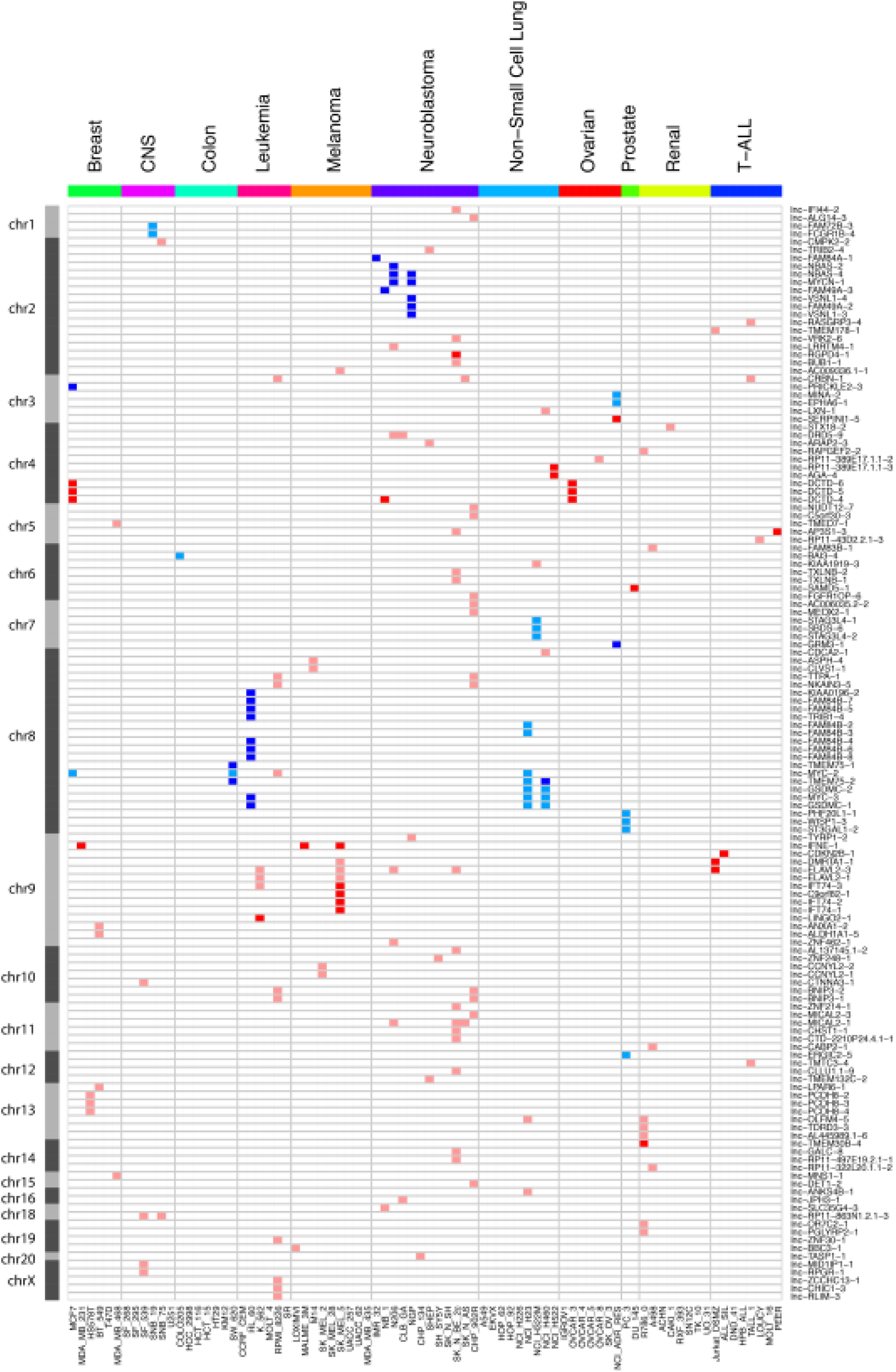
Overview of the IncRNA genes affected by focal SCNAs after extensive filtering. Red represents copy number loss (log-ratio < 1.5) in that cell line while blue corresponds to copy number gain (log-ratio > 1.5). Dark red and blue correspond to copy number changes with absolute log-ratio above 2.5.

### RT-qPCR confirms the majority of focal aberrations

We devised a unique strategy to validate the selected focal lncRNA SCNAs using qPCR. Assays were designed targeting the genomic locus of the aberration and the nearest exons of the flanking protein coding genes. By comparing the Cq value of the lncRNA locus and the flanking coding exons, we can accurately assess the difference in copy number between the two. Using this strategy, we evaluated 88 events (Figure 3). For 66 of these (75%) an altered copy number status compared to at least one of the two flanking assays could be confirmed, of which 43 (49%) showed the expected relative difference in Cq values with both flanking assays and were thus validated as focal aberrations. The validation rate is higher for the amplifications than for the deletions (56% and 48%, respectively). The validation rate drastically increases when we limit our analysis to the subset of segments with an absolute average log-ratio larger than 2.5. In that case, 58 out of 64 (91%) events are confirmed copy number alterations. The fraction of confirmed focal aberrations remains similar (53%).

**Figure 3.**
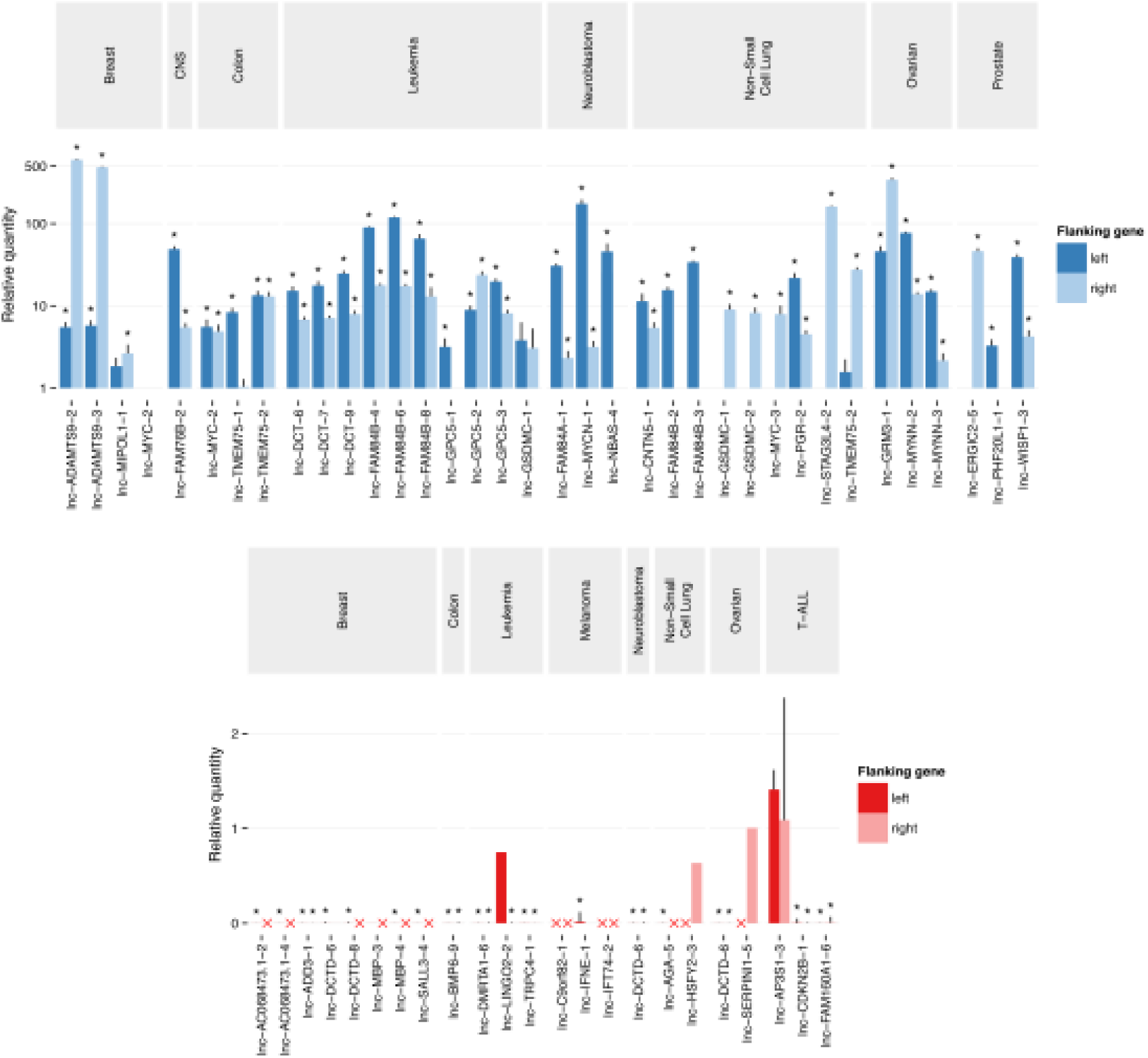
RT-qPCR validation of the putative focal SCNAs. The Cq value of the aberration is normalized to the Cq value of each of the flanking regions. A copy number gain (blue) is considered confirmed and focal when the relative quantity to both flanking regions is higher than 1. Similarly, a copy number loss (red) is considered confirmed and focal when the relative quantity to both flanking regions is less than 1. Red crosses represent Cq values > 35, corresponding to a homozygous deletion of the flanking regions. Stars represent significant (p-value < 0.05) differences from 1.

### Most novel lncRNA aberrations do not correspond to common somatic variants

As our custom platform differs considerably from other array CGH platforms, it not unlikely that the newly found SCNAs actually comprise uncharted germline copy-number variants that may exist in a normal population and do not contribute to cancer. To assess this possibility, we performed an RT-qPCR experiment for five validated loci on DNA from 192 healthy individuals. Neither homozygous deletions nor high order amplifications could be detected for any lncRNA in any of the samples (Figure S8). Of note, for one lncRNA heterozygous deletions were found in 12 individuals (6%).

## Discussion

Even though the number of samples we examined is limited and confined to cell lines, we were able to detect a large number of SCNA that specifically affect lncRNA exons. This suggests that similarly to protein-coding genes, lncRNAs are frequently targeted by SCNAs in cancer. After rigorous filtering focused on novel highly aberrant segments that not encompass protein coding genes, we report 136 such events, including 25 that are recurrent. Of those, 76 events were marked as focal based on the copy number of the flanking protein coding genes. Since the cancer genome harbors many large SCNAs, it is important to also consider the events where the flanking protein coding genes are not strictly copy number normal. As long as the lncRNA itself is focally affected by a second event as well.

Our strategy uncovered several cancer-associated lncRNAs. For instance, the known oncogene lnc-MYC-2 (PVT1) was detected as a recurrent focal aberration (Figure 2, Figure S3). PVT1 has been implicated in several cancer types including gastric cancer(22), ovarian cancer and breast cancer(18). PVT1 was found to be co-amplified in more than 98% of cancers with a MYC copy number increase(23). Our work not only confirms frequent amplification of PVT1 in cancer, but also reveals that PVT1 amplifications can be focal. Another interesting accordance with previous studies is found in a large-scale pan-cancer study on SCNAs(19). Although the authors mainly focus on SCNAs affecting protein coding genes and use limited lncRNA annotation, they report one lncRNA, lnc-DCTD-5 (LINC00290), as the sole member of a frequently deleted region. Our results reveal a recurrent and focal deletion in ovarian and breast cancer cell lines, suggesting a role in cancer (Figure 2).

The validation rate determined with qPCR was strongly dependent on the log-ratio cutoff applied to the segments, with an absolute average log-ratio larger than 2.5 showing high validation rates for lncRNA copy number status. The relatively high cutoff is likely to be related to the unique design of our platform. As the probes are confined to small genomic loci (lncRNA exons) is it not unimaginable that the observed signal-to-noise ratio is different compared to typical designs. In addition, qPCR may not be the most appropriate method to detect hemizygous copy number changes. Even with a stringent log-ratio cutoff (2.5), only 50% of the events could be confirmed to be truly focal. This suggests that the limited number of probes on the flanking protein coding genes is insufficient to define the breakpoints of the segments in some cases.

Nevertheless, even when taking the validation rate into account, our research finds about 100 lncRNAs affected by focal SCNA. As the majority of these events are likely no germline copy-number variants, these SCNAs harbor interesting candidates for further research.

## Conclusion

We developed and applied a unique array CGH platform capable of detecting small and focal lncRNA SCNAs. We have screened a panel of 80 cancer cell lines and shortlisted 136 IncRNA genes with a putative role in cancer. Among this list are several lncRNAs that have been implicated in cancer, validating our approach. Since the great majority of the lncRNAs on our platform have yet to be functionally studied, this finding suggests that our research provides many new cancer related lncRNA genes. We present a set of lncRNA genes to the lncRNA and cancer research community as novel candidate cancer lncRNA genes for further functional exploration.

## Methods

### LncRNA exon database

LncRNA transcript annotation was obtained from LNCipedia(24) (version 1.0) and stored in a MongoDB NoSQL database. Protein coding transcript annotation was obtained from Ensembl’s(25) biomart (version 64, September 2011) and stored in the same format. For every lncRNA transcript, the nearest upstream and downstream protein coding transcript was determined. To interface with the MongoDB dataset, both perl scripts and mongo shell scripts were employed. Using MongoDB’s MapReduce functionality, a non-redundant exon collection was built starting from the collection of non-redundant transcripts.

### Array CGH platform design

Array CGH probe design was performed using Agilent Technologies eArray softwarea. A BED file of all non-redundant exons was generated from the exon database and uploaded into eArray for probe design. Since our criterion to have 2 probes per exon was initially not met, the exon boundaries were extended and the corresponding BED files were uploaded as well. Exon boundaries are extended with 100 bp, 300 bp and 500 bp. In addition, less stringent selection parameters were used for the 500 bp extended exon. In this way, 5 probe datasets were generated and stored in a separate MongoDB collection. From this collection, 2 probes per exon (neighborhood) were selected with preference for the probes closest to the exon. Overlapping transcripts were taken into account to avoid duplicate probe selection. For transcripts with fewer than 5 exons, additional probes were selected until the transcript was covered by at least 10 probes. For the flanking protein coding genes, probes were designed for the 2 exons closest to the lncRNA. From this set, the 2 probes nearest to the lncRNA were selected. The resulting set of 166 417 unique probes was uploaded to eArray and supplemented with normalization and QC probe groups recommended by Agilent Technologies. Agilent Technologies subsequently manufactured the final design in the 4×180k format. The design of the platform is made publically available through the Gene Expression Omnibus (GEO) website using the accession number GPL22307.

### Cancer cell line DNA and RNA

The National Cancer Institute (NCI) provided DNA and RNA samples for all cell lines in the NCI 60 cancer cell line panel. The neuroblastoma and T-ALL cell lines were available in house; RNA extraction was performed with the miRNeasy Mini Kit (QIAGEN) and DNA extraction with the QIAamp DNA Mini Kit (QIAGEN).

### Array CGH

400 ng of genomic DNA was labeled with Cy3-dCTP (GE Healthcare, Belgium) using a Bioprime array CGH genomic labeling system (Invitrogen, Belgium). In parallel, Kreatech gender-matched controls were labeled with Cy5-dCTP. Samples were hybridized on the custom array CGH arrays for 40 h at 65 °C. After washing, the samples were scanned at 5 μm resolution using a DNA microarray scanner G2505B (Agilent Technologies). The scan images were analyzed using the feature extraction software 9.5.3.1 (Agilent Technologies). Segmentation is achieved using the circular binary segmentation algorithm in the DNACopy R package. Visual inspection and creation of the copy number profile plots is performed with ‘Vivar’(26). All raw array CGH data files are made publically available through the Gene Expression Omnibus (GEO) website using the accession number GSE85444.

### Segment analysis and filtering

Segment position and statistics are stored in a MongoDB collection. A perl script is used to combine the segment annotation with lncRNA and protein coding gene annotation in other collections and implement the filtering process. First, only segments that overlap lncRNA exons are retained. Next, segments with an absolute average log-ratio less than 1.5 are discarded as are segments contained within segmental duplications (UCSC genomicSuperDups track) or segments that overlap with more than 3 known variants (database of genomic variants(27)). The absolute log-ratio of the nearest segments covering the flanking protein coding genes should be 0.5 lower than the segment covering the lncRNA (corresponding to about 1 copy less). A more stringent subset of segments is obtained by requiring the absolute log-ratio of the nearest segments covering the flanking protein coding genes to be less than 0.35 (copy number neutral).

### RT-qPCR validation

QPCR assays are designed based on the chromosomal locations of the altered segment covering the lncRNA and the nearest exons of the two flanking protein coding genes. Primer design is performed using Primer3(28), primers spanning common SNPs are excluded. Specificity is evaluated using BiSearch(29). All qPCR reaction are prepared using Bio-Rad’s SsoAdvanced Universal SYBR Green Supermix in 5 μl (2.5 μL mastermix, 0.25 μl of each forward and reverse primers (250 nM final concentration) and 2 μl DNA (5 ng)). QPCR plates are analyzed on the LightCycler480 (Roche) using 2 min activation at 95 °C, followed by 45 cycles of 5 sec at 95 °C, 30 sec at 60 °C and 1 second at 72 °C, and a melt curve analysis.

Calculation of normalized relative quantities was done using the qbase+ software version 2.6 (Biogazelle) and the open source statistical environment R (version 3). The Cq values corresponding to the altered segment are normalized to those corresponding to the flanking protein coding genes and scaled to the control sample (Human Genomic DNA, Roche). Downstream analysis and data visualization was achieved using R and third party modules (plyr, ggplot2).

## Acknowledgments

This work was supported by the Multidisciplinary Research Partnership ‘Bioinformatics: From Nucleotides to Networks’ Project of Ghent University [01MR0310W to P.V]; Fund for Scientific Research Flanders [FWO; to P.M]

## Footnotes

a https://earray.chem.agilent.com/earray/

---

**Figure S1:**
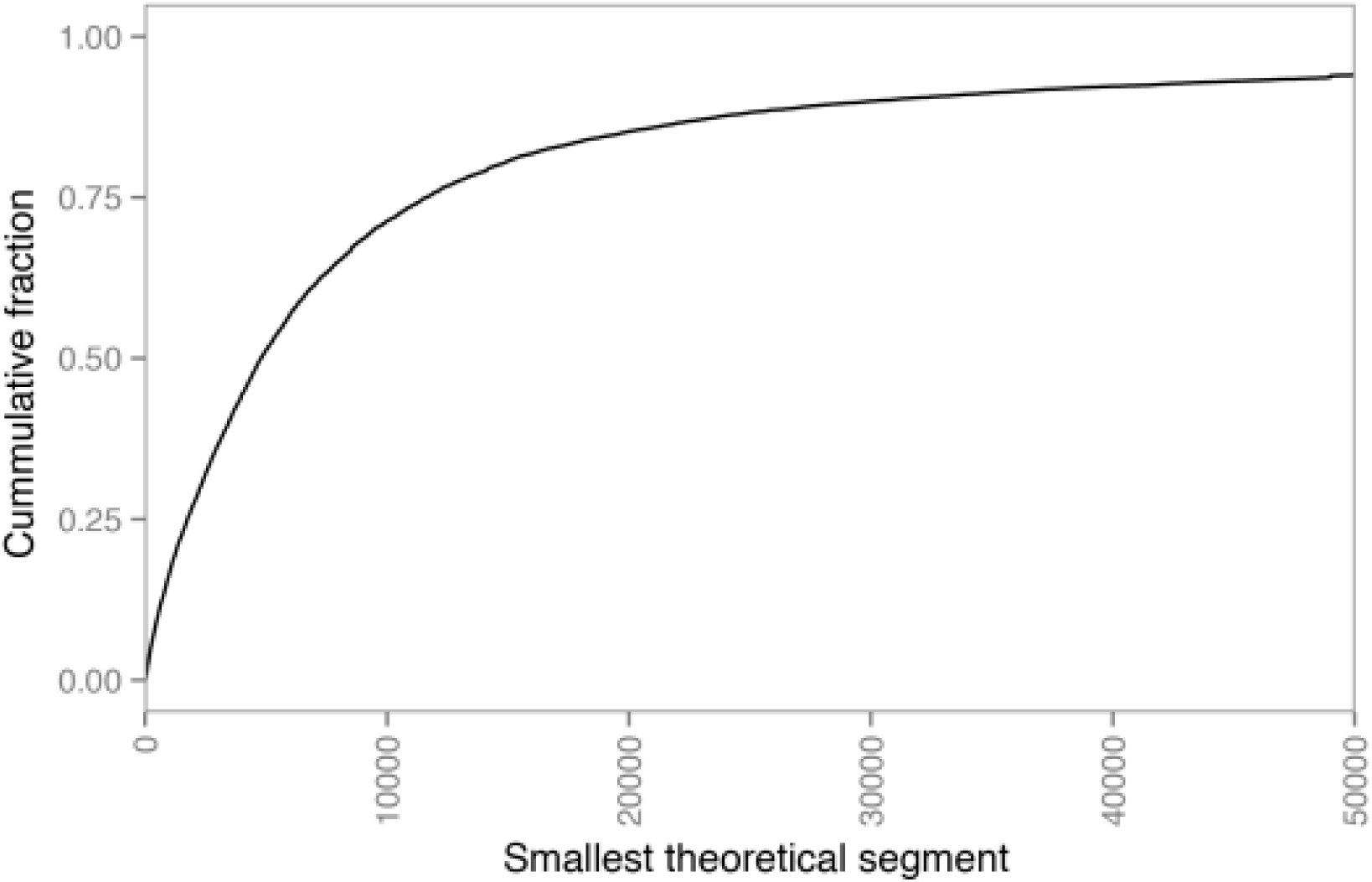
The smallest theoretical segment that covers each lncRNA using the Genome-Wide Human SNP Array 6.0. A theoretical segment is the distance between the two closest probes covering IncRNAs.

**Figure S2:**
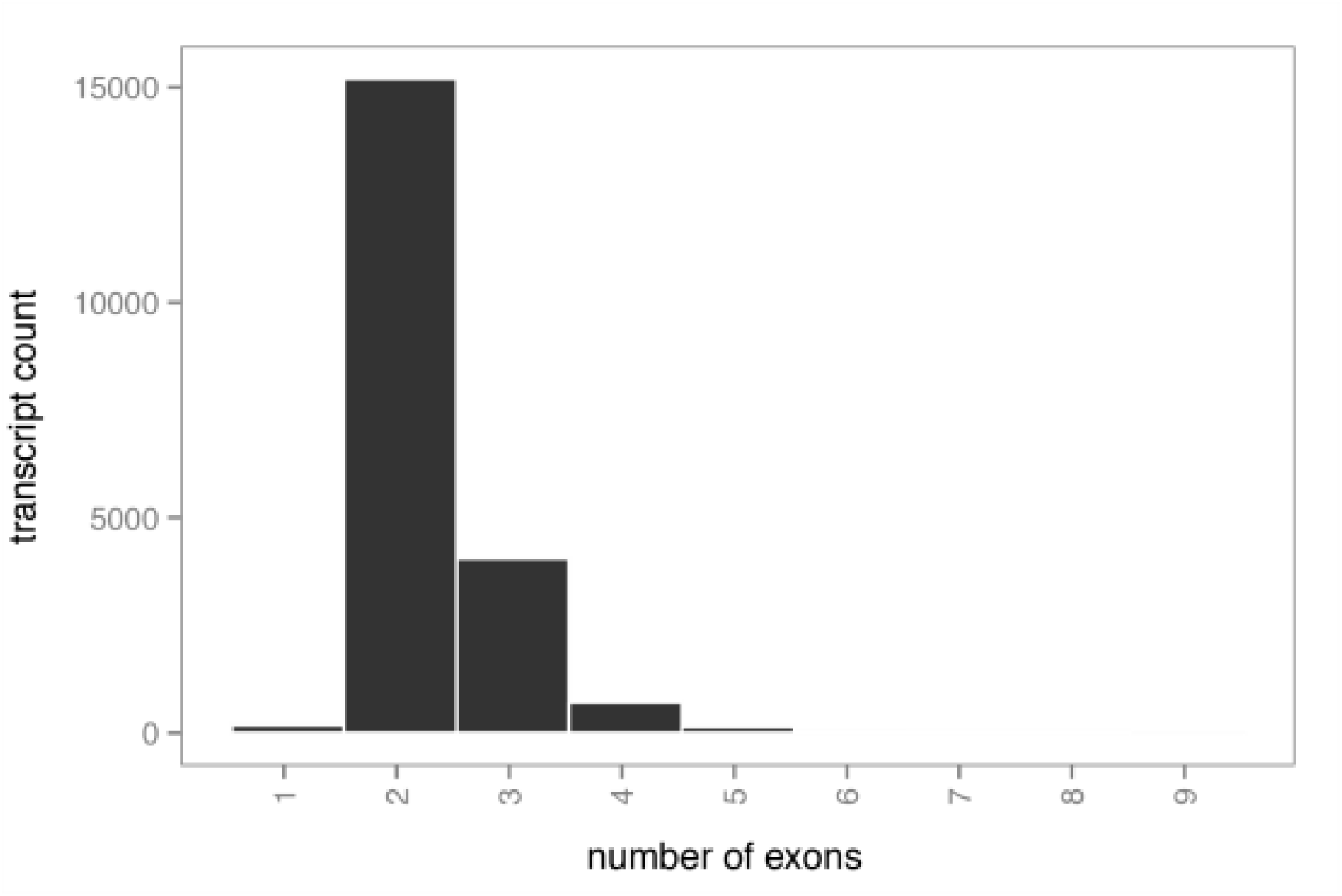
Distribution of the number of exons for the lncRNA transcripts in our dataset

**Figure S3:**
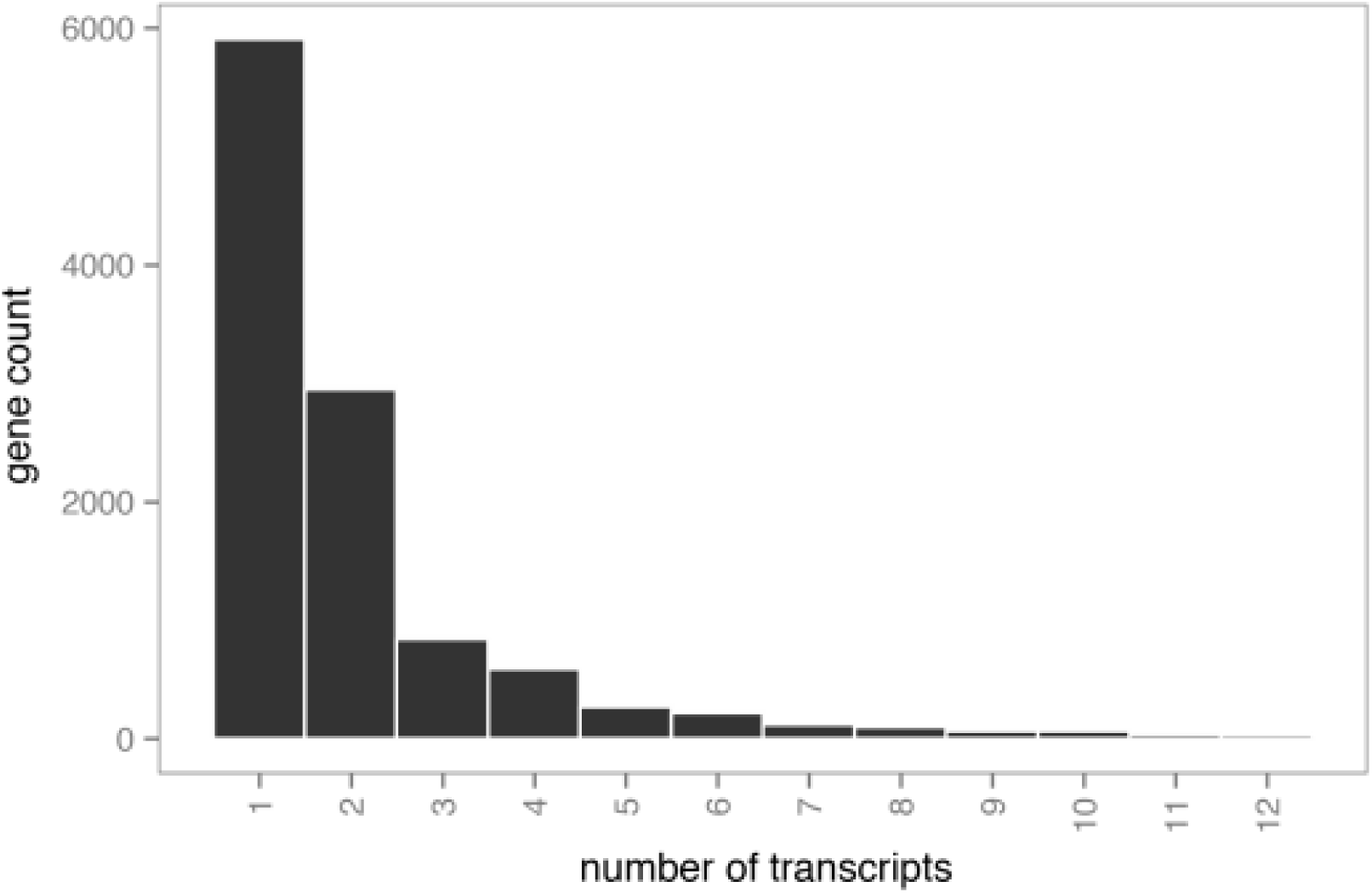
Distribution of the number of transcripts per lncRNA gene (locus).

**Figure S4:**
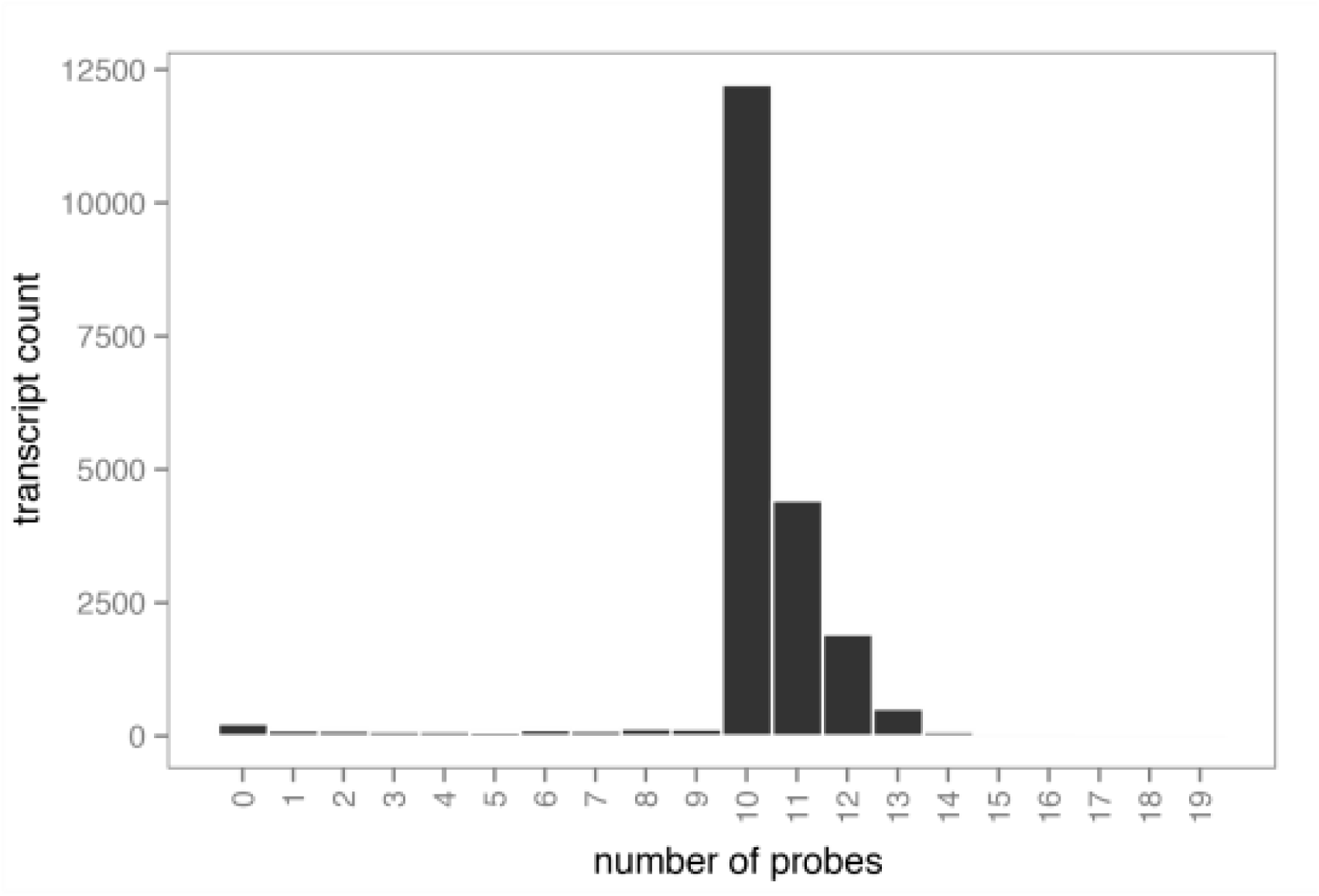
The required number of probes for a transcript depends on the number of exons. For transcripts with five exons or less, the required number of probes is 10. For larger transcripts, the required number is two times the number of exons. Only for a small fraction of transcripts, the pipeline failed to design at least 10 probes.

**Figure S5:**
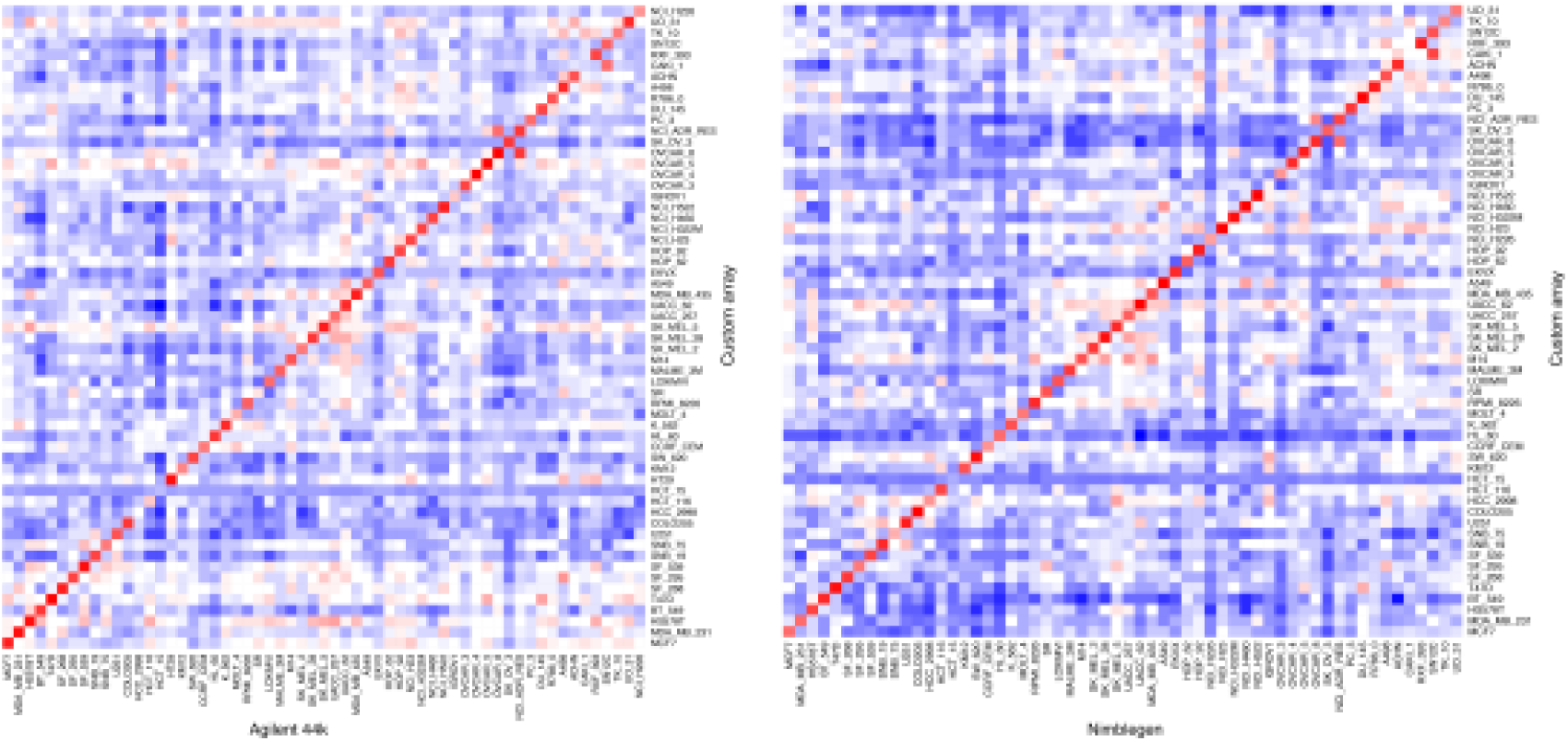
Comparison of the global copy number profiles (averaged in 1Mb bins) with publically available profiles of two different array CGH platforms (Agilent 44K and Nimblegen 385k). Pearson correlation of all samples is depicted. Blue corresponds to no correlation while red is a high correlation. Excellent correlation if observed for our platform. As expected, cell lines derived from the same individual (such as NCI/ADR-RES and OVCAR-8) are also highly correlated. In addition, this analysis revealed problems with 2 DNA samples (HCT-15 and CAKI-1) that were unresolved by repeating the hybridization.

**Figure S6:**
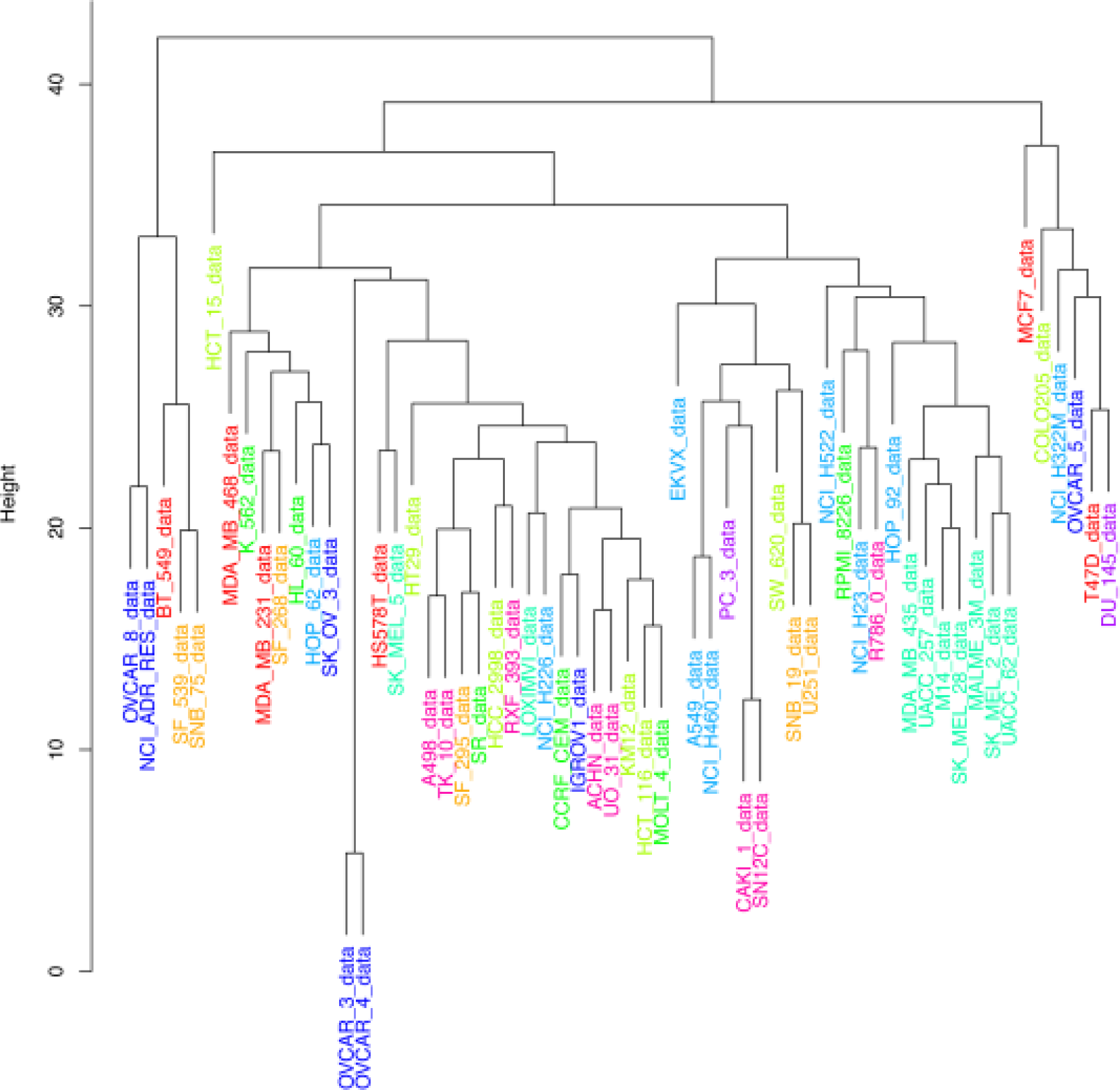
Complete linkage tree of the cell line based on their global copy number profiles. Cell lines derived from the same individual cluster closely together. On a cancer subtype level, only melanoma samples (green) cluster together.

**Figure S7:**
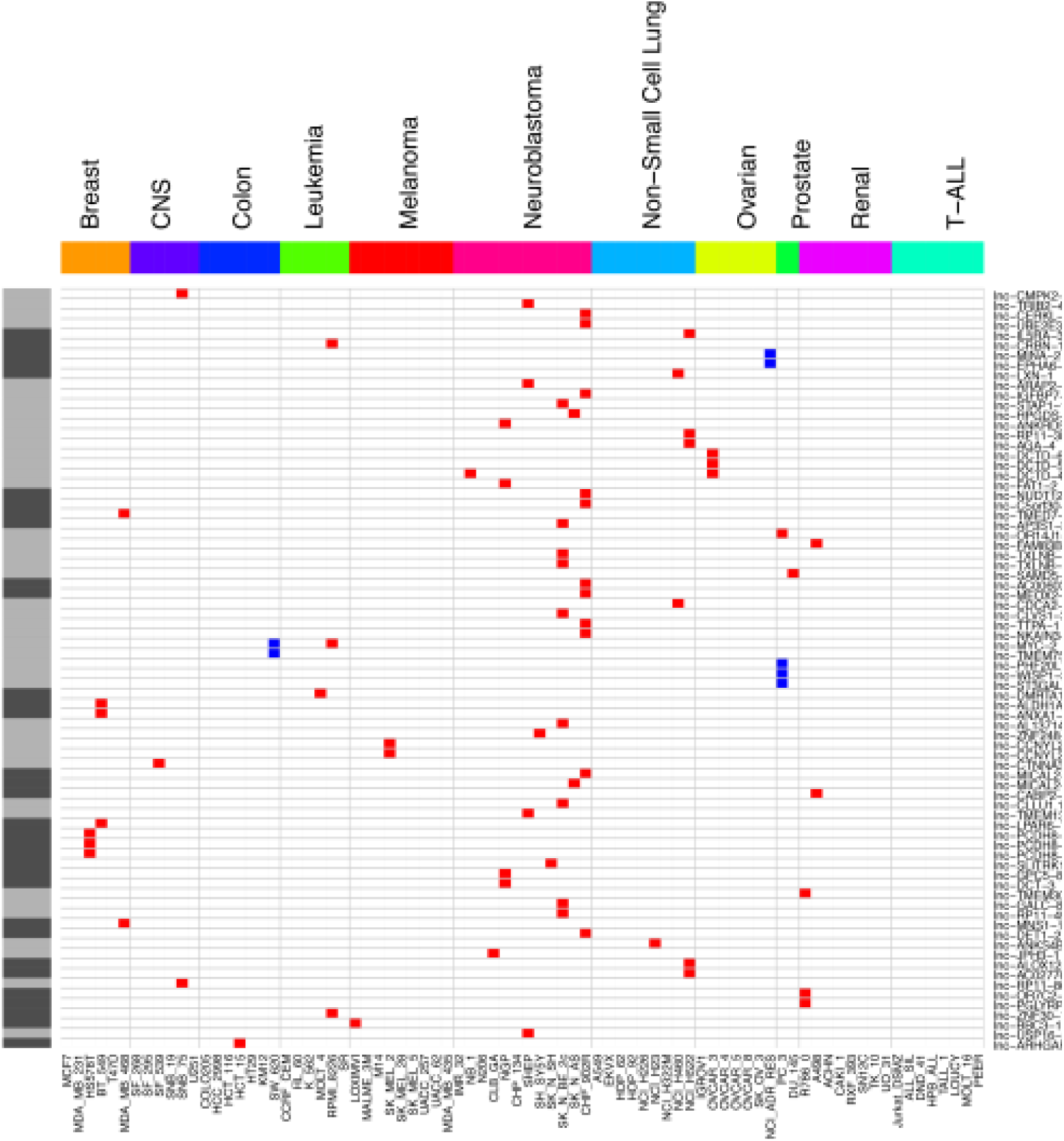
Overview of the IncRNA genes affected by focal SCNAs and copy number normal flanking protein coding genes. Red represents copy number loss (log-ratio < 1.5) in that cell line while blue corresponds to copy number gain (log-ratio > 1.5).

**Figure S8:**
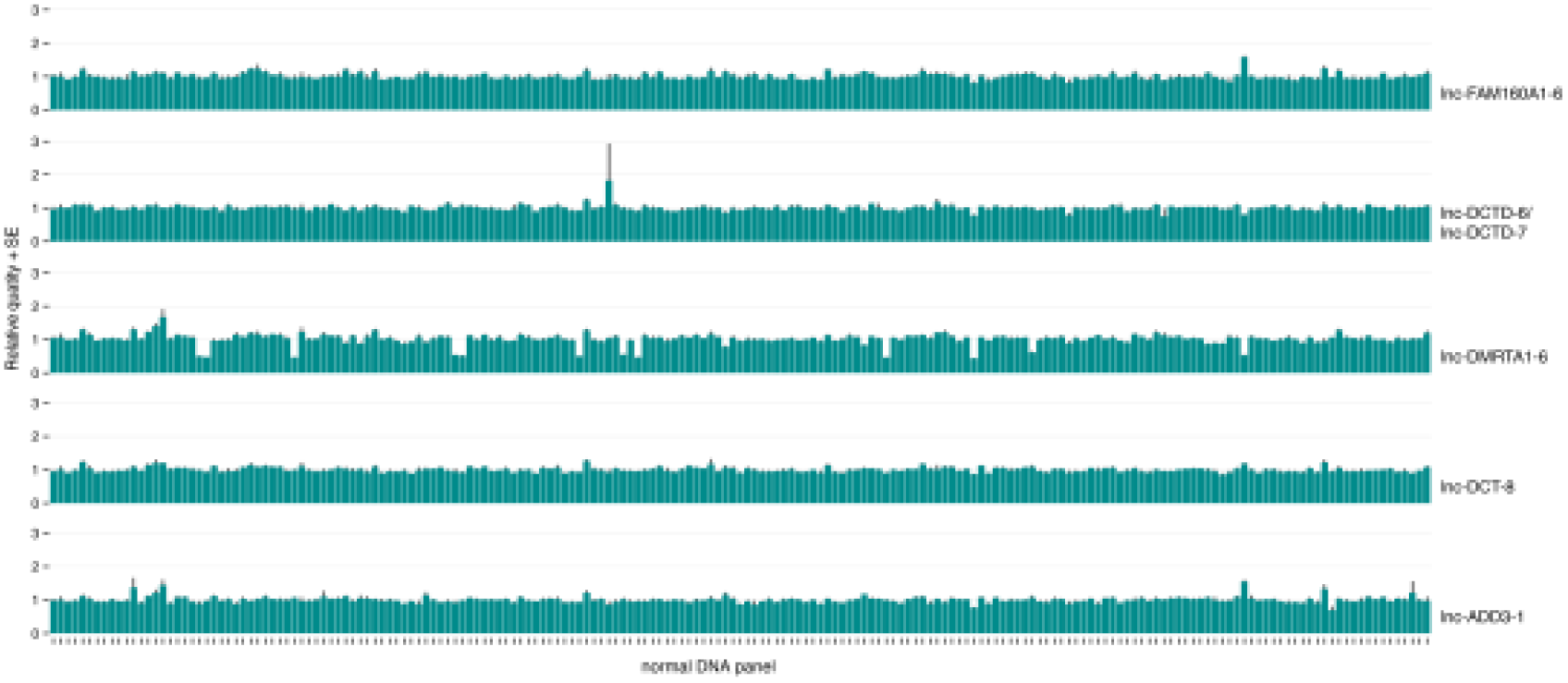
RT-qPCR validation of 5 selected loci on 192 DNA samples of healthy individuals. Neither homozygous deletions nor high order amplifications could be detected for any lncRNA in any of the samples. However, heterozygous lnc-DMRTA1-6 deletions were present in 12 samples (6%).

